# LuminoCell: a versatile and affordable luminometer platform for monitoring in-cell luciferase-based reporters

**DOI:** 10.1101/2022.01.28.478134

**Authors:** Kamila Weissová, Bohumil Fafílek, Tomasz Radaszkiewicz, Canan Celiker, Petra Macháčková, Tamara Čechová, Jana Šebestíková, Aleš Hampl, Vítězslav Bryja, Pavel Krejčí, Tomáš Bárta

## Abstract

Luciferase reporter assays represent a simple and sensitive experimental system in cell and molecular biology to study multiple biological processes. However, the application of these assays is often limited by the costs of conventional luminometer instruments and the versatility of their use in different experimental conditions. Therefore, we aimed to develop a small, affordable luminometer allowing continuous measurement of luciferase activity, designed for inclusion into various kinds of tissue culture incubators. Here we introduce LuminoCell - an open-source platform for the construction of an affordable, sensitive, and portable luminometer capable of realtime monitoring in-cell luciferase activity. The LuminoCell costs $40, requires less than 1 hour to assemble, and it is capable of performing real-time sensitive detection of both magnitude and duration of the activity of major signalling pathways in cell cultures, including receptor tyrosine kinases (EGF, FGF), WNT/β-catenin, and NF-κB. Additionally, we show that the LuminoCell is suitable to be used in cytotoxicity assays as well as for monitoring periodic circadian gene expression.

## Introduction

Luciferase reporter assays allow the study of a wide range of biological processes in cells and tissues. These reporter systems utilize luciferins, a class of small molecules that react with oxygen in the presence of the enzyme luciferase, to release the energy in the form of light. Luciferase assays represent a well-established experimental approach in cell and molecular biology, commonly used for gene expression analyses, promoter analyses, cytotoxicity assays, and signal transduction analyses. These assays are sensitive, reproducible, compatible with a range of internal control reporters, and provide a broad linear dynamic signal range (Smale, 2010). The extent of the application of luciferase assays in research is demonstrated by the number of publications that reference the luciferase reporters; over 47,000 research articles were found at Pubmed (keyword: “luciferase reporter”, December 2021), with more than 31,000 articles published in the past 10 years.

In luciferase reporter assays, the luciferase activity is determined by a luminometer device, usually in a bench microplate or a single tube reader configuration. However, high costs of commercial luminometers may limit their availability for many research groups. Additionally, there are other limitations associated with these devices: I) the endpoint measurement is often the only possible way of analysis and it does not provide information about the temporal dynamics of the signal; II) most instruments are difficult to sterilize and cannot be placed in the cell incubator, and thus are not suitable for long-term experiments with growing cell cultures; III) the measurement often requires cell harvest and lysis; IV) complex experimental schemes, such as multiple treatments at different times, are difficult to perform with the standard trap-door luminometer devices.

Here, we introduce an open-source platform for construction of a versatile, cheap, light-weight, and portable luminometer - the LuminoCell - that can be 3D printed and assembled within one hour. The LuminoCell enables continuous in-cell monitoring of luciferase activity in the growing cell cultures, allowing the study of dynamic biological processes in real-time. We also provide proof-of-concept experiments of the device use, in monitoring the activity of cell signalling pathways, cytotoxicity assays, and gene expression assays.

## Results

### The LuminoCell description

The central part of the LuminoCell is represented by the light to frequency converter TSL237S-LF (Mouser, USA) that is positioned in the centre of 3D printed wells in a plastic case (**Fig. 1A, B, C**). A light-to-frequency converter like the TSL237S-LF, converts light intensity into a series of square-wave pulses, with the frequency depending on the light intensity. The limit of light sensitivity is determined by the on-sensor noise, resulting in occasional spurious pulses even without incoming light (dark frequency, f_D_). The typical f_D_ of TSL237S-LF, according to the manufacturer, is 0.1 Hz and the maximum operating frequency is 1 MHz, providing a dynamic range of seven orders of magnitude per one second of the measurement process. Pulses, generated by TSL237S-LF upon light detection, are integrated by the Arduino microcontroller unit (MCU) (www.arduino.cc) that is positioned inside the LuminoCell. The luminometer is placed into a cell incubator and it is connected to a computer outside an incubator using a thin USB cable. The cell culture Petri dishes, containing luciferase reporter cell lines, are positioned on the device and covered by a lid to protect potential incoming light, while allowing gas exchange (**Fig. 1B, C**). Luciferase activity (number of pulses) is monitored using a serial monitor provided by Arduino IDE software (www.arduino.cc). The LuminoCell is capable of simultaneously measuring the luciferase signal in six 40 mm petri dishes and estimated costs for assembling the LuminoCell are ~40 USD (the costs for the 3D printed case are not included) (**Tab. S1**). The general wiring diagram of the LuminoCell is shown in **Fig. 1A** (details are depicted in **Fig. S1, S2**).

**Figure 1:**
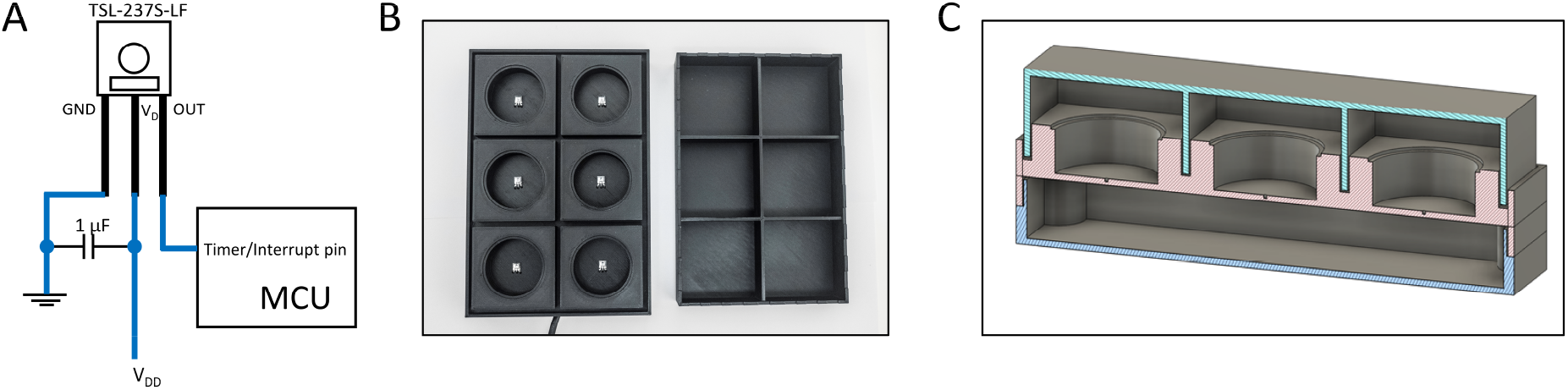
The LuminoCell description. (A) General wiring diagram. (B) The LuminoCell with lid (top view). (C) Cross section of the LuminoCell. green – lid, red – middle part, blue – bottom part.

### Proof-of-concept experiments

We aimed to test the performance of the LuminoCell using a wide spectrum of different luciferase reporters and cell lines. We tested the LuminoCell in monitoring the activity of fibroblast growth factor receptor (FGFR) signalling in cells, using the pKrox24^Luc^ luciferase reporter. The pKrox24 contains promoter sequences based on the *EGR1* upstream of the firefly luciferase, and was developed to record the FGF-mediated activation of the RAS-ERK MAP kinase pathway (Gudernova et al., 2017). <u>R</u>at <u>c</u>hondo<u>s</u>arcoma (RCS) cells, stably expressing the pKrox24^Luc^ (RCS::pKrox24^Luc^), were used to assess the signal background, defined as the number of pulses (dark frequency, f_D_), in the absence of luciferin. Six to seven pulses during a 5 minute integration time were recorded, which corresponds to f_D_=~0.02 Hz (**Fig. 2A**). When luciferin was added to the culture media, the number of recorded pulses increased to 15-18/5 minutes (f_D_=~0.06 Hz) reflecting the basal pKrox24^Luc^ transactivation. Addition of the recombinant FGFR ligand FGF2 led to a profound increase in the number of detected pulses, reaching its maximum at 142 pulses/5 minutes after 5 hours (300 minutes) of FGF2 addition (**Fig. 2A**). To remove the background, caused by the basal EGR1 activity in the absence of FGF2, we refined the data and subtracted the background noise (**Fig. 2B**). All the other data sets, shown in this manuscript, are presented with subtracted backgrounds. This experiment clearly demonstrates that the LuminoCell is capable not only to monitor kinetic changes of the luciferase activity in real-time, but also allows researchers to pick the right time for potential downstream analyses.

**Figure 2:**
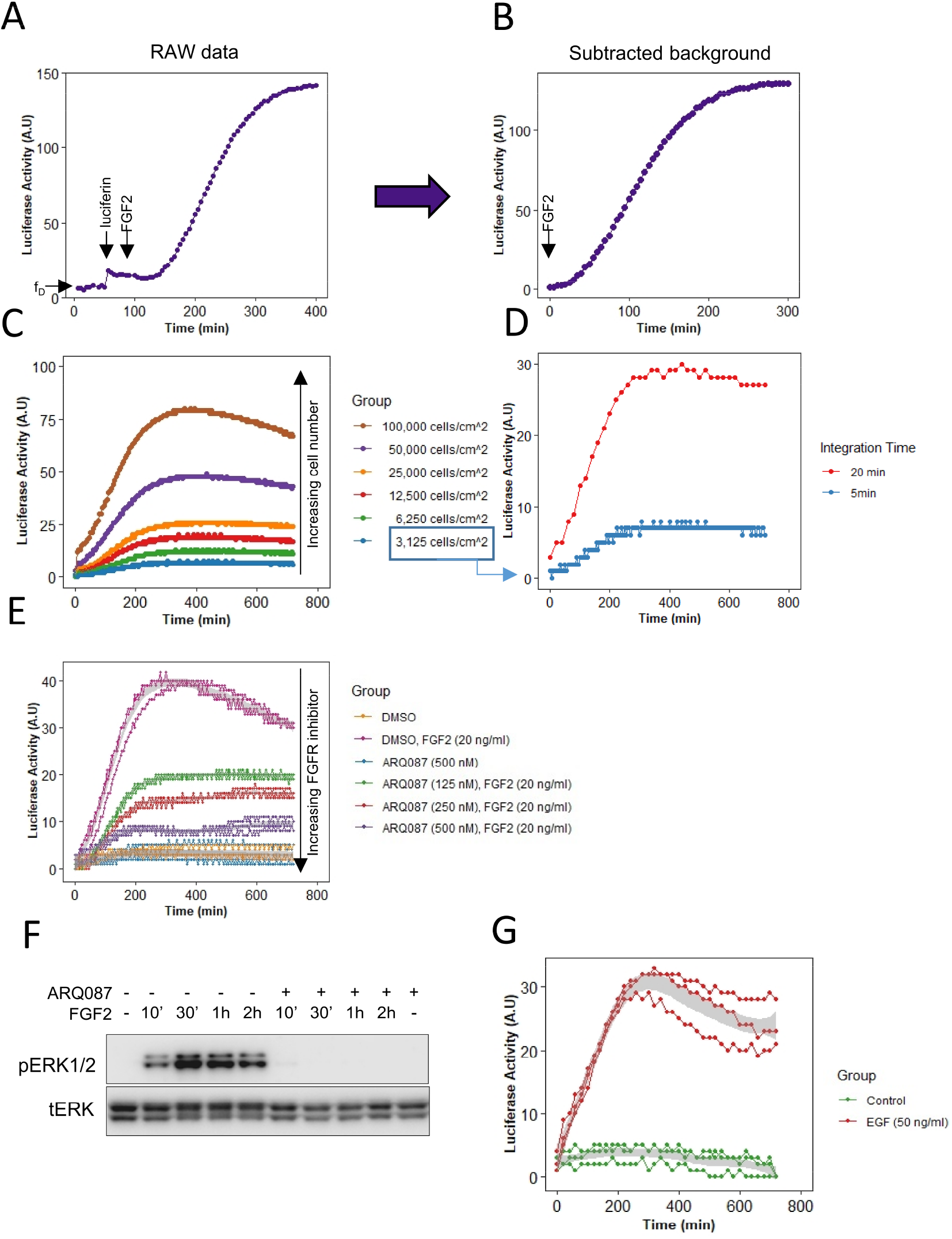
Transactivation of pKrox24^Luc^ by FGFR and EGFR signalling. (**A**) Luciferase activity upon addition of luciferin and recombinant FGF2 (arrows) to RCS::pKrox24^Luc^ cells. f_D_, background (dark) frequency. Integration time for pulse count was 5 minutes, if not stated otherwise. (**B**) Normalized data from (A), with subtracted background. (**C**) FGF2-mediated pKrox24^Luc^ transactivation in different cell densities. (**D**) 3,125 RCS::pKrox24^Luc^ cells/cm^2^ were stimulated with 20 ng/ml FGF2, and the luciferase activity was measured using a 5 or 20 minute integration time. (**E**) Luciferase activity in the presence of 20 ng/ml FGF2 and different concentrations of FGFR signalling inhibitor ARQ087 in RCS::pKrox24^Luc^ cells. (**F**) Western blot analysis of ERK phosphorylation (p) in RCS::pKrox24^Luc^ cells treated with 500 nM ARQ087 and 20 ng/ml FGF2. Total levels of ERK (t) serve as a loading control. (**G**) pKrox24^Luc^ transactivation mediated by EGF in 293T::pKrox24^Luc^ cells. Integration time 10 minutes. Data from three replicates are shown.

We evaluated different cell densities to test the sensitivity of signal detection. The RCS::pKrox24^Luc^ cells were seeded at concentrations ranging from 3×10^3^ to 1×10^5^ cells/cm^2^. The cells were stimulated with FGF2 and luciferase activity measurement started immediately upon FGF2 addition. The FGF2-mediated increase in signal was recorded even at the lowest used cell density (3×10^3^ cells/cm^2^), generating 8 pulses/5 minutes above the background (**Fig. 2C**). Extending the pulse integration time from 5 to 20 minutes led to an approximate 5 fold increase in luminometer sensitivity (8 vs. 30 pulses at 3×10^3^ cells/cm^2^), with a negligible effect on temporal resolution of the signal (**Fig. 2D**). Thus, the integration time may be varied to increase luminometer sensitivity, in order to detect the signal of weak promoters, dim luciferase reporters or a low number of cells.

To demonstrate the LuminoCell capacity to monitor dynamic changes of the reporter activity, we treated RCS::pKrox24^Luc^ cells with ARQ087, a potent inhibitor of FGFR catalytic activity (Balek et al., 2017). ARQ087 was added into the culture media 30 minutes prior to FGF2, with luciferase measurement startingimmediately upon the FGF2 addition (**Fig. 2E**). We found a substantial inhibition of the pKrox24^Luc^ transactivation in cells treated with ARQ087. This was confirmed by western blot analysis of FGF2-mediated ERK MAP kinase phosphorylation in RCS::pKrox24^Luc^ cells, which was inhibited by ARQ087 (**Fig. 2F, S3**).

The pKrox24^Luc^ responds to activation of the RAS-ERK module and therefore can be transactivated by many cell signalling pathways which use ERK, including as many as 30 different receptor tyrosine kinases (Gudernova et al., 2017). We tested whether the luminometer is able to monitor the activity of a receptor tyrosine kinase unrelated to FGFR, the epidermal growth factor receptor (EGFR). The 293T::pKrox24^Luc^ cells were cultured in the presence of EGF and luciferase activity was monitored. Figure 2G shows the pKrox24^Luc^ transactivation mediated by EGFR signalling.

We tested the activity of the canonical WNT/β-catenin pathway, using <u>S</u>uper-<u>T</u>op-<u>F</u>lash (STF) cells stably expressing luciferase reporter under the control of seven LEF/TCF binding sites (Xu et al., 2004). Cells were cultured in the presence or absence of GSK3α/β inhibitor CHIR99021, a potent activator of the canonical Wnt signalling cascade (Bennett et al., 2002). We detected an increase of luciferase signal 3 hours (180 minutes) upon CHIR99021 addition (**Fig. 3A**). The signal increased to 23-29 pulses after 10 hours (600 minutes), followed by a decrease to 10-15 pulses at the end of experiment, 16.5 hours (1000 minutes), while the control increased from 4 pulses at the start of measurement to 4-8 pulses at the end of experiment. End-point analysis of the Wnt signalling luciferase reporter activity using a bench-top luminometer confirmed activation of the Wnt signalling pathway 3 hours after CHIR99021 treatment (**Fig. 3B**). Additionally, western blot analysis confirmed the activation of the Wnt canonical pathway 3 hours after CHIR99021 treatment, demonstrated by accumulation of β-catenin that is associated with a decrease of its phosphorylation and followed by stabilization of its target gene *AXIN2* (**Fig. 3C, S4**).

**Figure 3:**
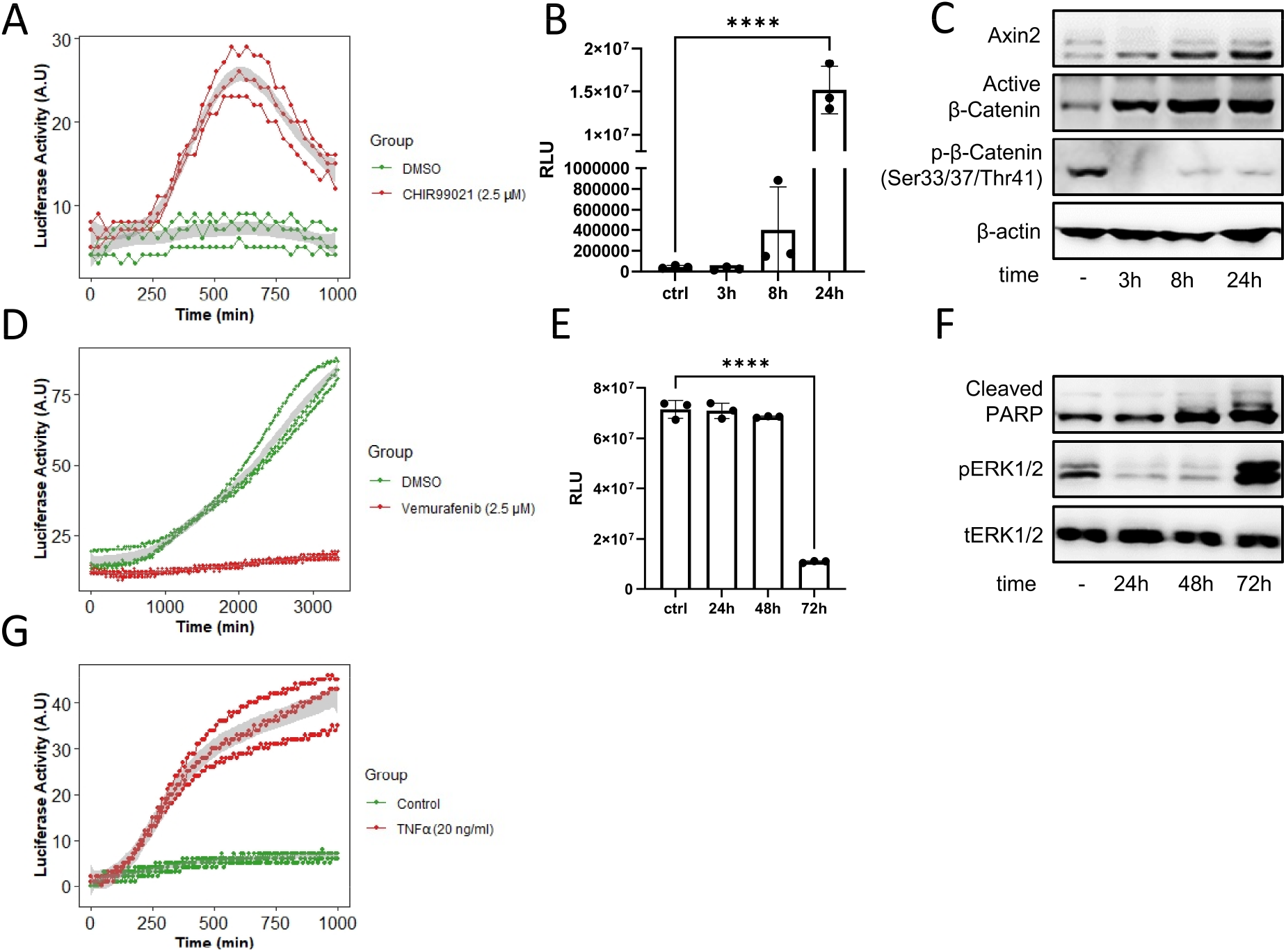
(**A**) Assessment of canonical Wnt signalling activity in STF cells in the presence of CHIR99021 or DMSO as a control. Luciferase activity measured using the LuminoCell (integration time 30 minutes). Data from three replicates are shown. (**B**) End-point analysis of canonical Wnt signalling activity in STF cells in the presence of CHIR99021 or DMSO as a control (ctrl). Luciferase activity measured using bench-top luminometer (Hidex Bioscan). (**C**) Western blot analysis of β-catenin and Axin2 expression in STF cells, β-actin represents a loading control. The data shows representative blots of three independent experiments. (**D**) Cytotoxicity testing using A375-IV^Luc^ cells. Cells were cultured in the presence of vemurafenib or DMSO as a control. Luciferase activity measured using the LuminoCell (integration time 20 minutes). Data from three replicates are shown. (**E**) Endpoint analysis of cytotoxicity using A375-IV^Luc^ cells. Cells were cultured in the presence of vemurafenib or DMSO as a control.). Luciferase activity measured using bench-top luminometer (Hidex Bioscan). (**F**) Western blot analysis of PARP cleavage and ERK (p) phosphorylation. Total (t) ERK represents a loading control. The data shows representative blots of three independent experiments. (**G**) Assessment of NF-κB activity in the presence of TNFα using 293T::NF-κB^Luc^ cells. Luciferase activity measured using the LuminoCell (integration time 10 minutes). Data from three replicates are shown.

Our experiments demonstrated the ability of the LuminoCell to monitor dynamic changes of the luciferase reporters for major signalling pathways. Next, we aimed to test the LuminoCell for its ability to be used for cell cytotoxicity assay. The melanoma cell line A375-IV^Luc^ (Kucerova et al., 2014) continuously expressing luciferase vector pGL4.50 (luc2/CMV/Hygro) (Promega) was cultured in the presence of Vemurafenib - an inhibitor of the constitutively active B-Raf V600E proto-oncogene, used for the treatment of late-stage melanoma (Chapman et al., 2011; Flaherty et al., 2010). Vemurafenib blocks cell proliferation, induces cellular stress and senescence of melanoma cells via inhibition of the B-Raf/MEK/ERK pathway (Haferkamp et al., 2013; Peng et al., 2017; Su et al., 2020). We detected an increase of luciferase activity in the control condition (DMSO) after 11 hours (660 minutes) from the start of the experiment that kept increasing to 87 pulses, indicating an increase of cell proliferation. However, in the presence of Vemurafenib we detected low luciferase activity ranging from 14 to 19 pulses at the end of the experiment, indicating blocked cell proliferation (**Fig. 3D**). End-point analysis using a bench-top luminometer revealed a decrease of luciferase activity 72 hours upon Vemurafenib treatment (**Fig. 3E**). Western blot analysis revealed increased apoptosis associated with increased PARP cleavage 48 hours after treatment and elevated phosphorylation of ERK (Su et al., 2020; Yue and López, 2020), indicating induction of cellular stress and paradoxical reactivation of the MAPK pathway in response to Vemurafenib treatment (Su et al., 2020) (**Fig. 3F, S5**).

We tested the activity of canonical NF-κB signalling in 293T cells stably expressing the NF-κB^Luc^ reporter (293T::NF-κB^Luc^). Cells were treated by recombinant TNFα that represents a potent activator of NF-κB (Schütze et al., 1992). The increasing intensity of the luciferase signal was recorded for at least ~16.5 hours (1000 minutes), generating up to 46 pulses/10 minutes (**Fig. 3G**). Cells in the absence of TNFα increased the luciferase signal from 0-2 pulses/10 minutes to 6-7 pulses/10 minutes at the end of the experiment at 16.5 hours.

We tested the LuminoCell for the ability to record circadian oscillations in human dermal fibroblasts (hDf). The use of luciferase reporters in circadian research highly simplifies and cuts the expenses for these long-term experiments, allowing the study of circadian rhythmic gene oscillation *in vitro* in real time. *BMAL1 (ARNTL)* is one of the key components of the circadian molecular clock and its expression exhibits a rhythmic pattern with an approximately daily period. We used luciferase *BMAL1* expression reporter and generated the hDf::BMAL1^Luc^ reporter cell line using the lentiviral transduction approach. Confluent hDf::BMAL1^Luc^ cultures were synchronized using serum shock, a widely-used approach to synchronize molecular circadian machinery (Balsalobre et al., 1998), and luciferase activity was monitored for 52 hours. Using the LuminoCell, we were able to detect high-amplitude circadian oscillations of *BMAL1* expression in hDFs cultures (**Fig. 4A**). This amplitude of *BMAL1* expression was confirmed using western blot analysis (**Fig. 4B**).

**Figure 4:**
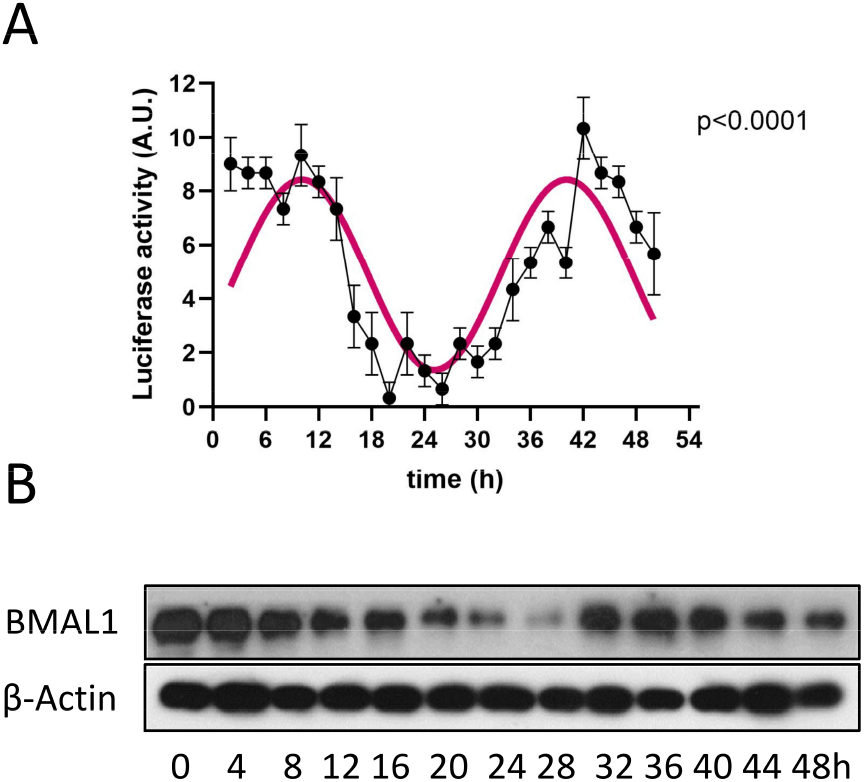
(**A**) Assessment of *BMAL1* expression using luciferase reporter cell line hDf::BMAL1^Luc^ upon serum shock (integration time 2h). Error bars show mean ±sd from three replicates. (**B**) Expression of BMAL1 upon serum shock, as demonstrated using western blot. β-actin represents loading control. The data shows representative blots of three independent experiments.

## Discussion

Here we introduce an open-source platform for construction of an affordable and simple LuminoCell, a luminometer capable of real-time monitoring of luciferase activity in living cells. We provide a full description of the device, 3D print files, and the complete source code shared via public repository. Additionally, we also provide a set of proof-of-concept experiments that test the LuminoCell functionality. Five different luciferase reporters expressing the firefly reporter gene *luc2* (*Photinus pyralis*) were used to test the luminometer. It is of note that the firefly luciferase belongs to a relatively weak reporter system, when compared to the *Renilla, Gaussia*, or NanoLuc systems, producing light emission centred at 560 nm (Gil et al., 2012). Given the broad detection range of different wavelengths of light (320-1050 nm), the LuminoCell is capable of detecting other light emission-based reporter systems, however one should test if the given reporter system is applicable to be used with this luminometer.

The LuminoCell provides a sufficient theoretical dynamic range of seven orders of magnitude (0.1 Hz-1 MHz), and it is comparable to expensive commercial devices that possess a dynamic range of about eight orders of magnitude. Although we have not compared the LuminoCell with other commercial devices, it is of note, however, that this luminometer cannot compete with commercial instruments in terms of detection limits. While this simple luminometer has a detection limit around 40 pW/s of light irradiance (TSL237S-LF has irradiance responsivity of 2.3 kHz/(μW/cm^2^)), devices equipped with expensive PMT tubes, or special CCD cameras offer much better sensitivity in attowatts (Enomoto et al., 2018). The relatively low detection limit of light irradiance is, however, estimated for 1 second of integration time. Therefore, one can improve the detection limits and dynamic range of this simple luminometer by increasing the integration time that is necessary to count pulses generated by TSL237S-LF, as described in **Fig. 2D, 4B**. However, this approach will lead to a decrease of temporal resolution, which is still sufficient to monitor dynamic changes of the luciferase activity during experiments.

Among the most beneficial advantages of the LuminoCell include: its price, versatility, small size, potential customization, and expandability. While the prices for commercial devices range from thousands to tens of thousands of USD, this luminometer provides a cheap and affordable solution for any research group. Additionally, commercial devices very often do not allow simultaneous real-time measurements in living cells and require end-point analysis often associated with cell lysis. The LuminoCell allows real-time measurements that do not require cell lysis, providing to a researcher the possibility to perform treatment of the reporter cell line during measurements and to continue to monitor dynamic changes of the luciferase activity upon treatment. Additionally, due to its compact size, the LuminoCell can be positioned into various kinds of tissue incubators including CO_2_ cell incubators, multi-gas incubators, hypoxia stations, and other specialized culture systems. Due to its portability, the LuminoCell can be used in a wide range of scientific disciplines ranging from cell and molecular biology to ecology for monitoring pollutants in the environment. Importantly, commercially available luminometers are not amenable for customization. The users are able to adapt, expand the potential, and customize the LuminoCell within open-source philosophy with forward compatibility by the scientific community. For example, one can expand the number of sensors to measure more samples simultaneously, or replace the TSL237S-LF with another sensor that would be better suited for the particular experimental system or reporter.

These abilities make the LuminoCell an applicable platform for a wide-range of experimental approaches that utilize luciferase reporter systems. Our work has enhanced the availability of the luminometer system with improved expandability and customizability, benefiting researchers from a wide range of life sciences disciplines.

## Material and Methods

### Cell culture and treatments

All cell lines were cultured in Knockout Dulbecco’s modified Eagle’s medium (DMEM), (Invitrogen, Life Technologies Ltd.) containing 10% foetal bovine serum (FBS), (PAA), 2 mM L-glutamine (Invitrogen, Life Technologies Ltd.), 1 × MEM nonessential amino acid solution, 1 × penicillin/streptomycin (PAA) and 10 μM β-mercaptoethanol (Sigma-Aldrich). The cells were incubated at 37 °C/5 % CO_2_ and regularly passaged using trypsin. For cell treatment the following inhibitors and growth factors were used: FGF (R&D Systems, 233-FB-025), EGF (Sigma Aldrich, SRP3196-500UG), TNF-α (R&D Systems, 210-TA-005), CHIR99021 (MedChem Express; HY-10182), Vemurafenib (MedChem Express; Hy-12057). Concentrations are indicated in each experiment.

### Luciferase vectors and reporter cell lines

For in-cell monitoring of the EGF and FGF signalling pathway activity, the 293T::pKrox24^Luc^ and RCS::pKrox24^Luc^ cells were produced from wt 293T and RCS cells using a piggyBac transposase for stable integration of TR01F plasmid (Mossine et al., 2013). NF-κB responsive element was replaced by Mapk-ERK responsive element from the pKrox24(MapErk)^Luc^ reporter (Gudernova et al., 2017). For assessment of the NF-κB pathway activity, the original TR01F was stably integrated into the 293T cells, again using the piggyBac transposase resulting in the 293T::NF-κB^Luc^ cells (Mossine et al., 2013). The clones with a successfully integrated cassette were selected as puromycin resistant and stimulated with appropriate ligands. Clones from each cell line with the best signal/noise ratio were used for the experiments. Plasmid pGL4.50(luc2/CMV/Hygro) (Promega) was used for generation of the A375-IV^Luc^ cell line by antibiotic selection (Hygromycin b, 200 μg/ml, Santa Cruz Biotechnology, 31282-04-9) and the limiting dilution method for clonal lines generation. SuperTOPFlash 293T (STF) cells were a kind gift from Q. Xu and J. Nanthas (Johns Hopkins University, Baltimore, MD, USA) (Xu et al., 2004). For *BMAIL1* expression monitoring, hDfs were transduced using lentiviral particles containing the pABpuro-BluF vector (pABpuro-BluF was a kind gift from Steven Brown (Addgene plasmid #46824; http://n2t.net/addgene:46824; RRID:Addgene_46824)) (Brown et al., 2005). Lentiviral particles were generated, as described in (Peskova et al., 2020, 2019). Upon transduction, hDf::BMAL1^Luc^ were cultured in the presence of Puromycin (0.7 μg/ml).

### Measuring luciferase activity

For continuous measurement using the LuminoCell: If not stated otherwise, 200,000 cells/well were seeded into a 40 mm cell culture Petri dish and the cells were allowed to grow for an additional 48 hours. Before each experiment, the cell culture medium was replaced with the medium of the same composition but without phenol red containing 100 μM D-Luciferin sodium salt (Merck, L6882). The source code (https://github.com/barta-lab/LuminoCell) was compiled using Arduino IDE software (v. 1.8.13) (www.arduino.cc) and uploaded into Arduino MCU using a USB cable. The luminometer was placed into a tissue incubator, the culture dishes were positioned into the luminometer and covered by a lid to protect the cells from potential incoming light. Luciferase activity was measured using a built-in serial monitor in the Arduino IDE software on a laptop computer Lenovo SL500 with an Ubuntu operating system (20.04 version). The measured values were saved into a CSV file and processed in R studio (version 1.3.1093). For serum shock treatment, the confluent culture of the hDf::BMAL1^Luc^ was kept for 48h in serum-free media, then treated with high-serum concentrated media (50%) for 2h, washed and replaced with serum-free (phenol free, 100 μM D-Luciferin sodium salt) media. For hDf::BMAL1^Luc^ data analysis, the data was analyzed using a single cosinor-based method (Cornelissen, 2014) with a constant period of 30 hours measured with the BioDare2 (Zielinski et al., 2014). The analysis was done in Prism 8 software (GraphPad, La Jolla, USA).

For end-point luciferase activity measurement using a bench-top luminometer: The STF cells were treated with DMSO or 2.5 μM CHIR99021 for 3h, 8h and 24h. The A375-IV^Luc^ cells were exposed to 2.5 μM Vemurafenib for 24h, 48h and 72h. After that time, the cells were washed with PBS, lysed, and subsequently an assay was performed accordingly to the manufacturer’s protocol (E1960; Promega) and luminescence was detected using the Hidex Bioscan Plate Chameleon Luminometer (Hidex).

### Western blot analysis

The cells were harvested into the sample buffer (125 mM Tris-HCl pH 6.8, 20% glycerol, 4% SDS, 5%β-mercaptoethanol, 0.02% bromophenol blue). The samples were resolved by SDS-PAGE, transferred onto a PVDF membrane, incubated with the primary antibodies (see the list of antibodies in **Table S2**) and with an anti-rabbit secondary antibody (Merck), and visualized by chemiluminescence substrate (Merck) using a Fusion Solo station (Vilber).

### Luminometer description and construction

Files for the 3D print, including the Arduino source code, are located in the GitHub repository (https://github.com/barta-lab/LuminoCell).

## Technical notes

### • The sensor

The TSL237S-LF sensor is a temperature compensated sensor which responds over the light range of 320 nm to 1050 nm. The sensor has an acceptable angle of 120° and therefore is positioned in the centre of a well, at a 13 mm distance from the sample (bottom of the cell culture dish) to gather light coming from the whole surface area of a dish (**Fig. S2**). We recommend adding a decoupling capacitor (at least 1 μF) between the GND and Vdd leading to a reduction of noise and potential voltage spikes.

### • Software interrupts and MCU

For pulse counting, we used the interrupt approach. When an external interrupt is triggered (in the case of a pulse generated by the sensor), the MCU will cease the operation of the running code and pass control to an associated interrupt service. Therefore, in the case of bright light coming into the sensor (e.g. a sunny day), the microcontroller may not proceed with the programme and the serial output displays no values. This has some implications for simultaneous reading from multiple sensors (wells). If the MCU receives too many pulses from one sensor, it cannot count pulses from the other sensor(s). However, this is not an issue for luciferase activity measurement, as the highest frequency of incoming pulses that we measured in this study was 0.47 Hz (142 pulses/5 minutes). Therefore, it is highly unlikely that in the case of luciferase measurement the incoming interrupts will prevent readings from other sensors.

As an MCU any Arduino developmental board can be used. We tested Arduino Nano Every, Arduino Nano, Arduino Micro, and Arduino Uno. However, the choice of MCU largely depends on the number of pins on the MCU capable of performing interrupt operations (for the details check the Arduino website https://www.arduino.cc/reference/en/language/functions/external-interrupts/attachinterrupt/).

### • 3D design

The LuminoCell consists of three parts that are printed on a conventional 3D printer: I) lid, II) middle part, and III) bottom part (**Fig. 1C, S2**). The lid contains 1 mm openings at its bottom rim that are embedded into a groove upon closing the lid. This prevents incoming light, while allowing gas and moisture exchange. The bottom and middle parts are assembled using four 3 mm self-tapping screws. The gap between the bottom and middle part is then sealed using silicon glue to prevent an increment of moisture inside the device.

### • Construction notes

Before you begin to assemble the LuminoCell, remove all indication LEDs from the MCU, as they may produce light that might interfere with the sensors (for the positions of the LEDs, check the MCU manual). Alternatively, place an MCU into a separate box outside a tissue incubator. In this case keep the wire(s) between MCU and sensors as short as possible. Secure the MCU on the bottom part using spacers (or any other preferred method). Connect the sensors with the MCU and capacitor using wires according to the scheme (**Fig. S1**). Connect the MCU with a USB cable. Assemble the bottom and middle part using 3 mm self-tapping screws. Cover the LuminoCell with the lid. Install Arduino IDE software (www.arduino.cc) onto your computer and run it. Connect the LuminoCell with a computer using the USB cable. Download the source code (.ino file) from https://github.com/barta-lab/LuminoCell and open it in Arduino IDE. Upload the code to the MCU and run the serial monitor (Ctrl+Shift+M). You should see the number of pulses that are detected in each well/sensor. The default integration time is set to 300,000 ms (5 minutes). If everything works and you plan to use the LuminoCell in a tissue incubator, seal the bottom part and middle part with silicon glue, additionally seal all holes in the wells for the sensor leads and opening for the USB cable.

## Acknowledgement

T. Barta is supported by the Czech Science Foundation (GA21-08182S) the Grant Agency of Masaryk University (GAMU) - MUNI/G/1391/2018. A. Hampl is supported by the European Regional Development Fund - Project INBIO (No.CZ.02.1.01/0.0/0.0/16_026/0008451). P. Krejci is supported by the Ministry of Education, Youth and Sports of the Czech Republic (LTAUSA19030), the Agency for Healthcare Research of the Czech Republic (NV18-08-00567) and the Czech Science Foundation (GA19-20123S, 21-26400K). P. Macháčková is supported by MEYS CR (LM2018129 Czech-BioImaging). V. Bryja is supported by the Czech Science Foundation (GX19-28347X).

## References

• Balek L, Gudernova I, Vesela I, Hampl M, Oralova V, Kunova Bosakova M, Varecha M, Nemec P, Hall T, Abbadessa G, Hatch N, Buchtova M, Krejci P. 2017. ARQ 087 inhibits FGFR signaling and rescues aberrant cell proliferation and differentiation in experimental models of craniosynostoses and chondrodysplasias caused by activating mutations in FGFR1, FGFR2 and FGFR3. Bone 105:57–66. doi:10.1016/j.bone.2017.08.016

• Balsalobre A, Damiola F, Schibler U. 1998. A serum shock induces circadian gene expression in mammalian tissue culture cells. Cell 93:929–937. doi:10.1016/s0092-8674(00)81199-x

• Bennett CN, Ross SE, Longo KA, Bajnok L, Hemati N, Johnson KW, Harrison SD, MacDougald OA. 2002. Regulation of Wnt signaling during adipogenesis. J Biol Chem 277:30998–31004. doi:10.1074/jbc.M204527200

• Brown SA, Fleury-Olela F, Nagoshi E, Hauser C, Juge C, Meier CA, Chicheportiche R, Dayer J-M, Albrecht U, Schibler U. 2005. The period length of fibroblast circadian gene expression varies widely among human individuals. PLoS Biol 3:e338. doi:10.1371/journal.pbio.0030338

• Chapman PB, Hauschild A, Robert C, Haanen JB, Ascierto P, Larkin J, Dummer R, Garbe C, Testori A, Maio M, Hogg D, Lorigan P, Lebbe C, Jouary T, Schadendorf D, Ribas A, O’Day SJ, Sosman JA, Kirkwood JM, Eggermont AMM, Dreno B, Nolop K, Li J, Nelson B, Hou J, Lee RJ, Flaherty KT, McArthur GA, BRIM-3 Study Group. 2011. Improved survival with vemurafenib in melanoma with BRAF V600E mutation. N Engl J Med 364:2507–2516. doi:10.1056/NEJMoa1103782

• Cornelissen G. 2014. Cosinor-based rhythmometry. Theor Biol Med Model 11:16. doi:10.1186/1742-4682-11-16

• Enomoto T, Kubota H, Mori K, Shimogawara M, Yoshita M, Ohmiya Y, Akiyama H. 2018. Absolute bioluminescence imaging at the single-cell level with a light signal at the Attowatt level. BioTechniques 64:270–274. doi:10.2144/btn-2018-0043

• Flaherty KT, Puzanov I, Kim KB, Ribas A, McArthur GA, Sosman JA, O’Dwyer PJ, Lee RJ, Grippo JF, Nolop K, Chapman PB. 2010. Inhibition of mutated, activated BRAF in metastatic melanoma. N Engl J Med 363:809–819. doi:10.1056/NEJMoa1002011

• Gil JS, Machado HB, Herschman HR. 2012. A method to rapidly and accurately compare the relative efficacies of non-invasive imaging reporter genes in a mouse model and its application to luciferase reporters. Mol Imaging Biol 14:462–471. doi:10.1007/s11307-011-0515-1

• Gudernova I, Foldynova-Trantirkova S, Ghannamova BE, Fafilek B, Varecha M, Balek L, Hruba E, Jonatova L, Jelinkova I, Kunova Bosakova M, Trantirek L, Mayer J, Krejci P. 2017. One reporter for in-cell activity profiling of majority of protein kinase oncogenes. eLife 6:e21536. doi:10.7554/eLife.21536

• Haferkamp S, Borst A, Adam C, Becker TM, Motschenbacher S, Windhövel S, Hufnagel AL, Houben R, Meierjohann S. 2013. Vemurafenib Induces Senescence Features in Melanoma Cells. J Invest Dermatol 133:1601–1609. doi:10.1038/jid.2013.6

• Kucerova L, Skolekova S, Demkova L, Bohovic R, Matuskova M. 2014. Long-term efficiency of mesenchymal stromal cell-mediated CD-MSC/5FC therapy in human melanoma xenograft model. Gene Ther 21:874–887. doi:10.1038/gt.2014.66

• Mossine VV, Waters JK, Hannink M, Mawhinney TP. 2013. piggyBac transposon plus insulators overcome epigenetic silencing to provide for stable signaling pathway reporter cell lines. PloS One 8:e85494. doi:10.1371/journal.pone.0085494

• Peng U, Wang Z, Pei S, Ou Y, Hu P, Liu W, Song J. 2017. ACY-1215 accelerates vemurafenib induced cell death of BRAF-mutant melanoma cells via induction of ER stress and inhibition of ERK activation. Oncol Rep 37:1270–1276. doi:10.3892/or.2016.5340

• Peskova L, Cerna K, Oppelt J, Mraz M, Barta T. 2019. Oct4-mediated reprogramming induces embryonic-like microRNA expression signatures in human fibroblasts. Sci Rep 9:15759. doi:10.1038/s41598-019-52294-3

• Peskova L, Jurcikova D, Vanova T, Krivanek J, Capandova M, Sramkova Z, Sebestikova J, Kolouskova M, Kotasova H, Streit L, Barta T. 2020. miR-183/96/182 cluster is an important morphogenetic factor targeting PAX6 expression in differentiating human retinal organoids. Stem Cells Dayt Ohio. doi:10.1002/stem.3272

• Schütze S, Potthoff K, Machleidt T, Berkovic D, Wiegmann K, Krönke M. 1992. TNF activates NF-kappa B by phosphatidylcholine-specific phospholipase C-induced “acidic” sphingomyelin breakdown. Cell 71:765–776. doi:10.1016/0092-8674(92)90553-o

• Smale ST. 2010. Luciferase assay. Cold Spring Harb Protoc 2010:pdb.prot5421. doi:10.1101/pdb.prot5421

• Su Y, Ko ME, Cheng H, Zhu R, Xue M, Wang J, Lee JW, Frankiw L, Xu A, Wong S, Robert L, Takata K, Yuan D, Lu Y, Huang S, Ribas A, Levine R, Nolan GP, Wei W, Plevritis SK, Li G, Baltimore D, Heath JR. 2020. Multi-omic single-cell snapshots reveal multiple independent trajectories to drug tolerance in a melanoma cell line. Nat Commun 11:2345. doi:10.1038/s41467-020-15956-9

• Xu Q, Wang Y, Dabdoub A, Smallwood PM, Williams J, Woods C, Kelley MW, Jiang L, Tasman W, Zhang K, Nathans J. 2004. Vascular development in the retina and inner ear: control by Norrin and Frizzled-4, a high-affinity ligand-receptor pair. Cell 116:883–895. doi:10.1016/s0092-8674(04)00216-8

• Yue J, López JM. 2020. Understanding MAPK Signaling Pathways in Apoptosis. Int J Mol Sci 21:E2346. doi:10.3390/ijms21072346

• Zielinski T, Moore AM, Troup E, Halliday KJ, Millar AJ. 2014. Strengths and limitations of period estimation methods for circadian data. PloS One 9:e96462. doi:10.1371/journal.pone.0096462

