## Supplementary information for "LuminoCell: a versatile and affordable luminometer platform for monitoring in-cell luciferase-based reporters"

Supplementary Figures

**Supplementary Figure 1**

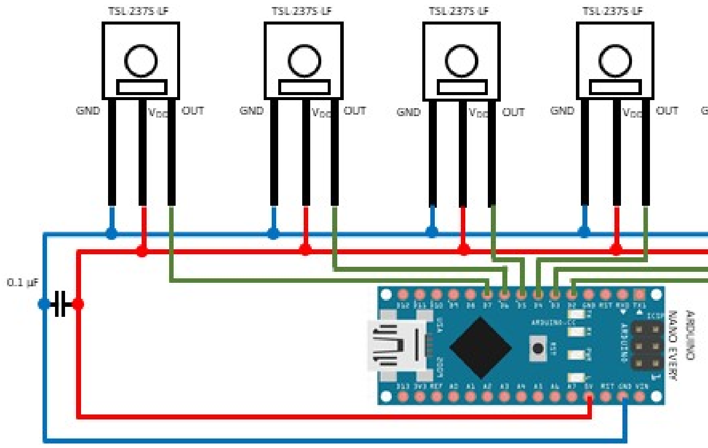


**Figure S1:** Wiring diagram of the LuminoCell device

**Supplementary Figure 2**

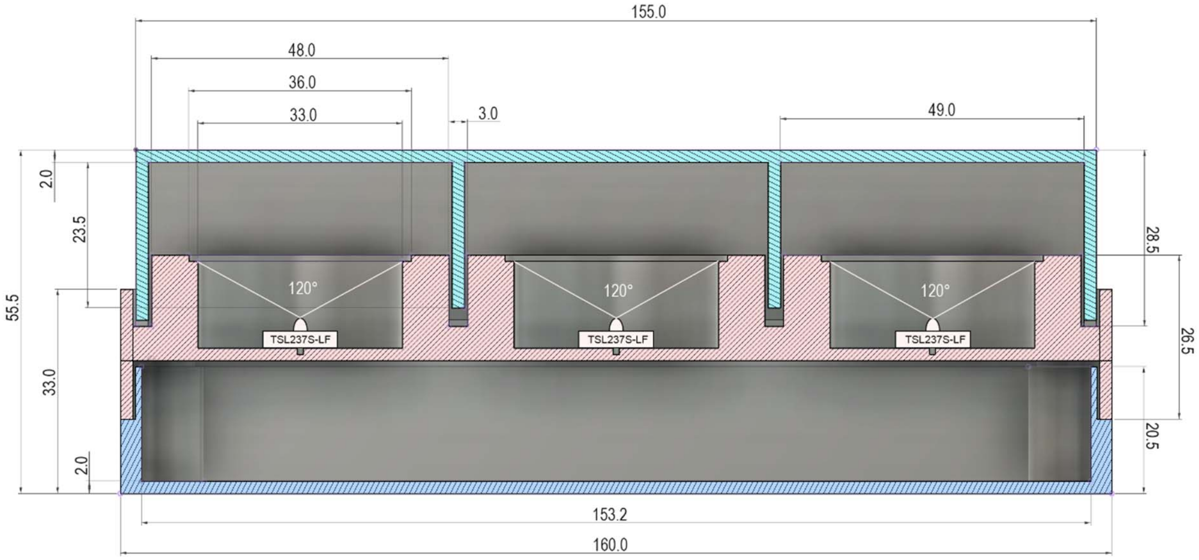


**Figure S2:** Cross section of the LuminoCell. Dimensions are given in millimetres.

**Supplementary Figure 3**

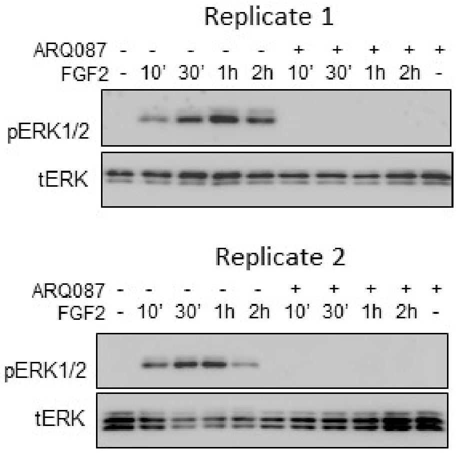


**Figure S3:** Western blot analysis of ERK phosphorylation (p) in RCS::pKrox24[Luc] cells treated with 500 nM ARQ087 and 20 ng/ml FGF2. Total levels of ERK (t) represent a loading control.

**Supplementary Figure 4**

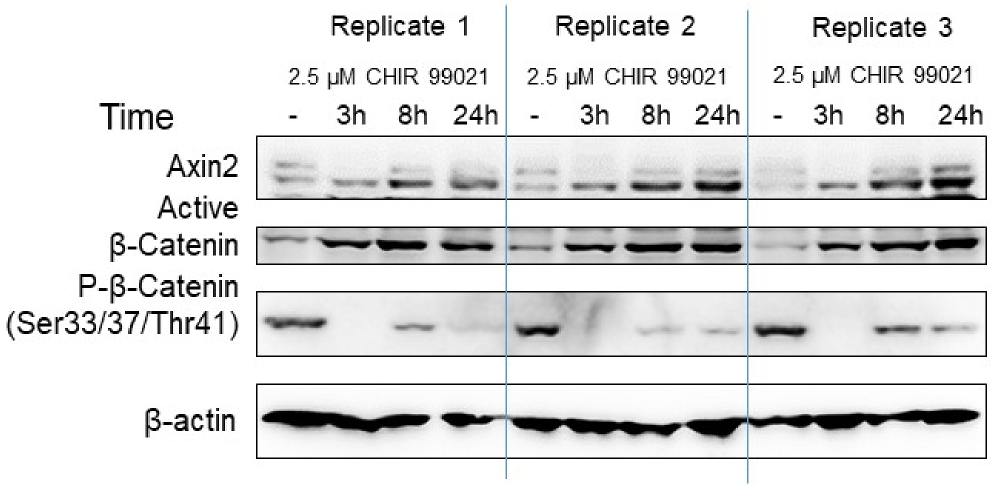


**Figure S4:** Western blot analysis of β-catenin and Axin2 expression in STF cells, β-actin represents a loading control.

**Supplementary Figure 5**

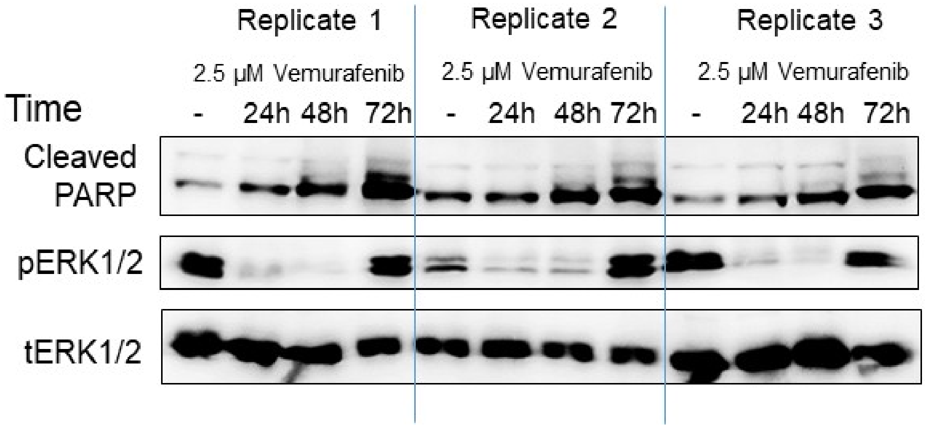


**Figure S5:** Western blot analysis of PARP cleavage and ERK (p) phosphorylation. Total (t) ERK represents a loading control.

**Supplementary Figure 6**

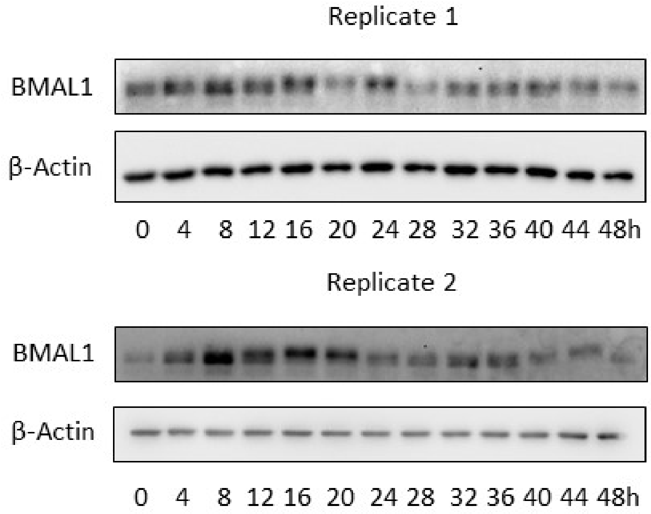


**Figure S6:** Western blot analysis of BMAL1 expression in hDF upon serum shock treatment, β-actin represents a loading control.

### Supplementary Tables

#### Supplementary Table 1

| Costs to build the LuminoCell | | |
|---|---|---|
| Quantity | Part | Price (USD) |
| 6x | TSL237S-LF | 24 |
| 1x | Arduino Nano Every | 11 |
| 1x | Capacitor (0.1 μF) | 0.1 |
| 1x | USB cable | 4 |
| **Estimated total costs** | | **39.1** |

**Table S1:** Summary of costs to build the LuminoCell. Costs for 3D printed case are not included.

#### Supplementary Table 2

| **List of primary antibodies used for Western blot analysis** | | |
|---|---|---|
| Name | Catalogue Number | Manufacturer |
| Axin2 | 2151 | Cell Signaling Technology |
| β-Catenin | 05-665 | Merck |
| Phospho-β-Catenin (Ser33/37/Thr41) | 9561 | Cell Signaling Technology |
| β-Actin | 4970 | Cell Signaling Technology |
| Cleaved PARP | 9541 | Cell Signaling Technology |
| pERK1/2 | 9101 | Cell Signaling Technology |
| tERK1/2 | 4695 | Cell Signaling Technology |
| BMAL1 | ab93806 | Abcam |

**Table S2:** Summary of primary antibodies used in this study.
